# Neutralization against Omicron SARS-CoV-2 from previous non-Omicron infection

**DOI:** 10.1101/2021.12.20.473584

**Authors:** Jing Zou, Hongjie Xia, Xuping Xie, Chaitanya Kurhade, Rafael R. G. Machado, Scott C. Weaver, Ping Ren, Pei-Yong Shi

**Author notes:** Correspondence: P.R. or P.-Y.S. J.Z., H.X., and X.X. contributed equally to this study.

## Abstract

The explosive spread of the Omicron SARS-CoV-2 variant underscores the importance of analyzing the cross-protection from previous non-Omicron infection. We developed a high-throughput neutralization assay for Omicron SARS-CoV-2 by engineering the Omicron spike gene into an mNeonGreen USA-WA1/2020 SARS-CoV-2 (isolated in January 2020). Using this assay, we determined the neutralization titers of patient sera collected at 1- or 6-months after infection with non-Omicron SARS-CoV-2. From 1- to 6-month post-infection, the neutralization titers against USA-WA1/2020 decreased from 601 to 142 (a 4.2-fold reduction), while the neutralization titers against Omicron-spike SARS-CoV-2 remained low at 38 and 32, respectively. Thus, at 1- and 6-months after non-Omicron SARS-CoV-2 infection, the neutralization titers against Omicron were 15.8- and 4.4-fold lower than those against USA-WA1/2020, respectively. The low cross-neutralization against Omicron from previous non-Omicron infection supports vaccination of formerly infected individuals to mitigate the health impact of the ongoing Omicron surge.

## Main

Severe acute respiratory syndrome coronavirus 2 (SARS-CoV-2) continues to evolve, leading to the emergence of variants of concern (VoC), variants of interest, and variants of monitoring. These variants can increase viral transmission, immune evasion, and/or disease severity.^1–3^ The recently emerged Omicron variant (B.1.1.529) was first identified in South Africa on November 2, 2021, and was designated as a new VoC on November 26, along with the four previous VoCs: Alpha, Beta, Gamma, and Delta.^4^ Since its emergence, Omicron has rapidly spread to over 89 countries, with case doubling in as little as 1.5 to 3 days, leading to global surges of COVID-19 cases.^5^ Compared to prior variants, the Omicron spike glycoprotein has accumulated more spike mutations, with over 34 mutations, many of which are known to evade antibody neutralization (*e.g.*, K417N, N440K, S477N, E484A and Q493R) or to enhance spike/hACE2 receptor binding (*e.g.*, Q498R, N501Y, and D614G).^1,3,6,7^ The high number of spike mutations is associated with decreased potency of antibody therapy and increased breakthrough Omicron infections in vaccinated and previously infected individuals^5^. Laboratory studies are urgently needed to examine the susceptibility of Omicron SARS-CoV-2 to vaccine- and infection-elicited neutralization. This study aimed to examine the cross-neutralization of Omicron by antibodies derived from previous non-Omicron infection.

To measure neutralization of the Omicron variant, we developed a high-throughput assay. Using a previously established mNeonGreen (mNG) reporter USA-WA1/2020 SARS-CoV-2,^8^ we swapped the original spike gene with an Omicron spike (BA.1 lineage; GISAID EPI_ISL_6640916), resulting in recombinant mNG Omicron-spike SARS-CoV-2 (**Extended Data Fig. 1**). The mNG gene was placed at the open-reading-frame-7 (ORF7) of the viral genome.^9^ The engineered Omicron spike contained mutations A67V, H69-V70 deletion (Δ69-70), T95I, G142D, V143-Y145 deletion (Δ143-145), N211 deletion (Δ211), L212I, L214 insertion EPE (Ins214EPE), G339D, S371L, S373P, S375F, K417N, N440K, G446S, S477N, T478K, E484A, Q493R, G496S, Q498R, N501Y, Y505H, T547K, D614G, H655Y, N679K, P681H, N764K, D796Y, N856K, Q954H, N969K, and L981F (**Extended Data Fig. 1**). The mNG Omicron-spike virus was sequenced to ensure no undesired mutations. After infecting Vero E6 cells, parental mNG USA-WA1/2020 developed larger fluorescent foci than Omicron-spike SARS-CoV-2 (**Extended Data Fig. 2**); however, comparable infectious titers of >10^6^ focus-forming units per milliliter (FFU/ml) were obtained for both viruses. The mNG viruses were used to develop a fluorescent focus reduction neutralization test (FFRNT) as depicted in **Extended Data Fig. 3**.

We examined the cross-neutralization of non-Omicron SARS-CoV-2-infected patient sera against Omicron virus. Two panels of COVID-19 patient sera, one collected at 1-month post-infection (n=64) and another collected at 6-month post-infection (N=36), were measured for their 50% fluorescent focus reduction neutralization titers (FFRNT_50_, defined as the maximal dilution that neutralized 50% of foci) against both USA-WA1/2020 and Omicron-spike SARS-CoV-2. **Extended Data Tables 1** and **2** summarize the patient information (*e.g.*, age, gender, race, date of positive viral test, symptom, and hospitalization) for the 1- and 6-month post-infection serum panels. All patients were infected before February 2021, prior to the emergence of the Omicron variant. The 1-month post-infection sera neutralized USA-WA1/2020 and Omicron-spike SARS-CoV-2 with geometric mean titers (GMTs) of 601 and 38, respectively (**Fig. 1a** and **Extended Data Fig. 4a**). Only one serum had a neutralization titer of <20 against USA-WA1/2020, whereas 23 of 64 sera had neutralization titers of <20 against Omicron-spike SARS-CoV-2 (**Fig. 1a**). Sera with high neutralization titers against USA-WA1/2020 were from symptomatic patients, most were hospitalized (**Extended Data Table 1**), confirming that neutralizing antibody levels are associated with COVID-19 disease severity.^10^ Notably, many of the sera with the highest neutralization titers of >3,450 against USA-WA1/2020 were from patients who had received convalescent plasma treatment (**Extended Data Table 1**).

**Figure 1.**
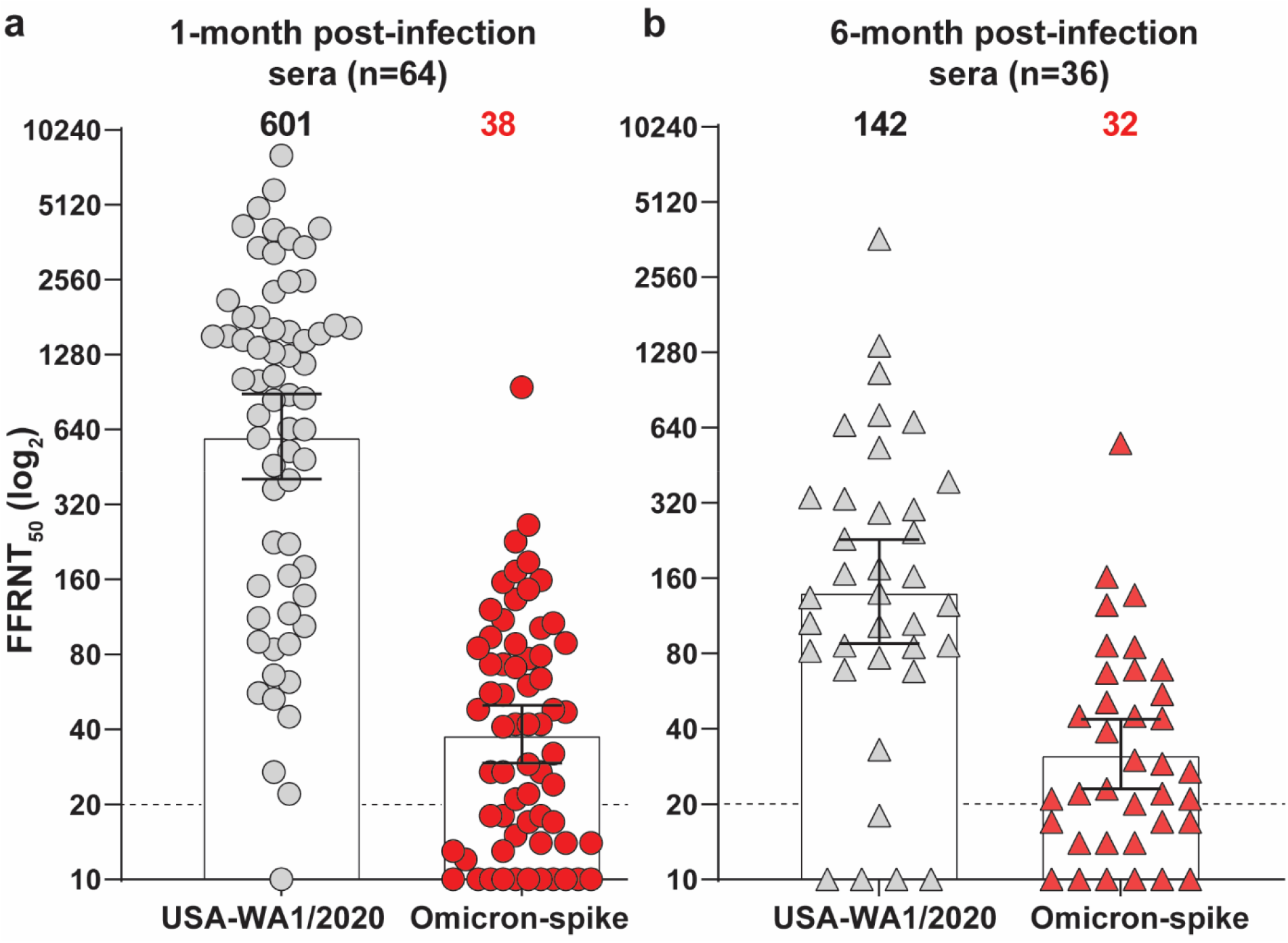
Reduced neutralization of Omicron SARS-CoV-2 by previous non-Omicron viral infection. 50% fluorescent focus reduction neutralization titers (FFRNT_50_) were measured for two serum panels from patients previously infected with non-Omicron SARS-CoV-2. The first serum panel was collected at 1-month post-infection (n=64) and the second panel collected at 6-months post-infection (n=36). For each serum, FFRNT_50_ values were determined against mNG USA-WA1/2020 and Omicron-spike SARS-CoV-2. **a,** FFRNT_50_s of 1-month post-infection sera. **b,** FFRNT_50_s of 6-month post-infection sera. **Extended Data Tables 1** and **2** summarize the FFRNT_50_ values and serum information for (**a**) and (**b**), respectively. Each symbol of dots (**a**) and triangles (**b**) represents one serum specimen. The FFRNT_50_ value for each serum was determined in duplicate assays and is presented as the geometric mean. The bar heights and the numbers above each set of data indicate geometric mean titers. The whiskers indicate 95% confidence intervals. The dotted line indicates the first serum dilution (1:20) of the FFRNT assay. The FFRNT_50_ values of sera that did not show any inhibition of viral infection are presented as 10 for plot purposes and statistical analysis. Statistical analysis was performed using the Wilcoxon matched-pairs signed-rank test. The statistical significance of the difference between the geometric mean titers against USA-WA1/2020 and Omicron-spike SARS-CoV-2 is p <0.0001 in both (**a**) and (**b**).

The 6-month post-infection sera neutralized USA-WA1/2020 and Omicron-spike SARS-CoV-2 with GMTs of 142 and 32, respectively (**Fig. 1b and Extended Data Fig. 4b**). Consistent with the 1-month post-infection results, symptomatic hospitalized patients tended to have higher neutralization titers against USA-WA1/2020 than asymptomatic individuals (**Extended Data Table 2**). Thus, from 1- to 6-months post-infection, the mean neutralization titers against USA-WA1/2020 waned from 601 to 142 (a 4.2-fold decrease), while the neutralization titers against Omicron-spike virus remained low and nearly unchanged at 38 and 32, respectively. Consistent with our results, the waning neutralization overtime against non-Omicron SARS-CoV-2 was previously reported in naturally infected or vaccinated individuals.^11–13^ Our data showed that 1- and 6-months after non-Omicron SARS-CoV-2 infections, the neutralization titers against Omicron were 15.8- and 4.4-fold lower than those against USA-WA1/2020, respectively. A similar range of neutralization reduction against the Omicron virus was reported for two-dose mRNA-vaccinated sera.^14–16^ Collectively, these results demonstrate low cross-neutralization against the Omicron variant from previous non-Omicron viral infection or two-dose mRNA vaccination. The low cross-neutralization against the Omicron variant strongly suggests that individuals previously infected with SARS-CoV-2 should be vaccinated to mitigate Omicron-mediated infection, disease, and potential death.

Among all tested sera, only 6 pairs of 1- and 6-month samples were collected from same individuals (**Extended Data Tables 1** and **2**). Their neutralization patterns (**Extended Data Fig. 5**) were similar to those observed with the means from complete 1- and 6-month serum panels.

Our study has several limitations. First, we have not defined the contributions of individual spike mutations to Omicron neutralization evasion. The constellation of Omicron spike mutations may result from selection for either viral transmission, immune escape, or both. The emergence of the Omicron variant in South Africa, where herd immunity is believed to be high, is consistent with evolutionary pressure for immune escape as suggested by our data and others.^17^ Second, the genotypes of viruses that infected the patients whose sera were analyzed in this study were not defined, although the timing suggests the Alpha variant. Third, we have not analyzed other immune modulators. CD8^+^ T cells and non-neutralizing antibodies that can mediate antibody-dependent cytotoxicity are known to protect patients from severe disease. The Omicron spike mutations may not dramatically affect T cell epitopes.^18^

The rapid evolution of SARS-CoV-2 underscores the importance of surveillance for new variants and their impact on viral transmission, disease severity, and immune evasion. Surveillance, laboratory investigation, and real-world vaccine effectiveness are essential to guide if and when an Omicron-specific vaccine or booster is needed. Currently, vaccination with booster shots,^19,20^ together with masking and social distance, remain to be the most effective means to mitigate the health impact of Omicron surge. Finally, the high-throughput fluorescent neutralization assay reported in this study can expedite therapeutic antibody screening, neutralization testing, and modified vaccine development.

## Methods

### Construction of recombinant viruses

The recombinant mNeoGreen (mNG) Omicron-spike SARS-CoV-2 was constructed on the genetic background of an infectious cDNA clone derived from clinical strain WA1 (2019-nCoV/USA_WA1/2020) containing an *mNG* reporter gene.^9^ The Omicron spike mutations, including A67V, Δ69-70, T95I, G142D, Δ143-145, Δ211, L212I, Ins214EPE, G339D, S371L, S373P, S375F, K417N, N440K, G446S, S477N, T478K, E484A, Q493R, G496S, Q498R, N501Y, Y505H, T547K, D614G, H655Y, N679K, P681H, N764K, D796Y, N856K, Q954H, N969K, and L981F, were engineered using a PCR-based mutagenesis protocol as reported previously.^1^ The full-length genomic cDNAs were *in vitro* ligated and used as templates to transcribe full-length viral RNA. Mutant viruses were recovered on day 3 after Vero E6 cells were electroporated with the *in vitro* RNA transcripts. The harvested virus stocks were quantified for their infectious titers (fluorescent focus units) by titrating the viruseson Vero E6 cells in a 96-well plate after 16 h of infection. The genome sequences of the virus stocks were confirmed to have no undesired mutations by Sanger sequencing. The detailed protocol of genome sequencing was recently reported.^21^

### Serum specimens

The research protocol regarding the use of human serum specimens was reviewed and approved by the University of Texas Medical Branch (UTMB) Institutional Review Board (IRB#: 20-0070). The de-identified convalescent sera from COVID-19 patients (confirmed by the molecular tests with FDA’s Emergency Use Authorization) were heat-inactivated at 56°C for 30 min before testing.

### Fluorescent focus reduction neutralization test

Neutralization titers of human sera were measured by a fluorescent focus reduction neutralization test (FFRNT) using the mNG reporter SARS-CoV-2. Briefly, Vero E6 cells (2.5 × 10^4^) were seeded in each well of black μCLEAR flat-bottom 96-well plate (Greiner Bio-one™). The cells were incubated overnight at 37°C with 5% CO_2_. On the following day, each serum was 2-fold serially diluted in the culture medium with the first dilution of 1:20. The diluted serum was incubated with 100-150 fluorescent focus units (FFU) of mNG SARS-CoV-2 at 37°C for 1 h (final dilution range of 1:20 to 1:20,480), after which the serum-virus mixtures were inoculated onto the pre-seeded Vero E6 cell monolayer in 96-well plates. After 1 h infection, the inoculum was removed and 100 μl of overlay medium (DMEM supplemented with 0.8% methylcellulose, 2% FBS, and 1% P/S) was added to each well. After incubating the plates at 37°C for 16 h, raw images of mNG fluorescent foci were acquired using Cytation™ 7 (BioTek) armed with 2.5× objective and processed using the default software setting. The foci in each well were counted and normalized to the non-serum-treated controls to calculate the relative infectivities. The curves of the relative infectivity versus the serum dilutions (log_10_ values) were plotted using Prism 9 (GraphPad). A nonlinear regression method was used to determine the dilution fold that neutralized 50% of mNG SARS-CoV-2 (defined as FFRNT_50_). Each serum was tested in duplicates.

## Statistics

The nonparametric Wilcoxon matched-pairs signed rank test was used to analyze the statistical significance in **Figure 1**.

## Data availability

The data that support the findings of this study are available from the corresponding authors upon request.

## Acknowledgments

We thank colleagues at Pfizer and the University of Texas Medical Branch for helpful discussion. P.-Y.S. was supported by NIH grants HHSN272201600013C, AI134907, AI145617, and UL1TR001439, and awards from the Sealy & Smith Foundation, the Kleberg Foundation, the John S. Dunn Foundation, the Amon G. Carter Foundation, the Gilson Longenbaugh Foundation, and the Summerfield Robert Foundation. S.C.W. was supported by NIH grant R24 AI120942.

## Author contributions

Conceptualization, X.X., P.R., P.-Y.S.; Methodology, J.Z., H.X., X.X., C.K., R.R.G.M., S.C.W., P.R., P.-Y.S; Investigation, J.Z., H.X., X.X., C.K., R.R.G.M., S.C.W., P.R., P.-Y.S; Resources, H.X., S.C.W., P.R., P.-Y.S; Data Curation, J.Z., H.X., X.X., P.R., P.-Y.S.; Writing-Original Draft, J.Z., H.X., X.X., R.R., P.-Y.S; Writing-Review & Editing, X.X., S.C.W., P.R., P.-Y.S.; Supervision, X.X., S.C.W., P.R., P.-Y.S.; Funding Acquisition, S.C.W., P.-Y.S.

## Ethics declarations Competing interests

The authors declare competing interests. X.X. and P.-Y.S. have filed a patent on the reverse genetic system. J.Z., H.X., X.X., and P.-Y.S. received compensation from Pfizer for COVID-19 vaccine development. Other authors declare no competing interests.

---

**Extended Data Figure 1.**
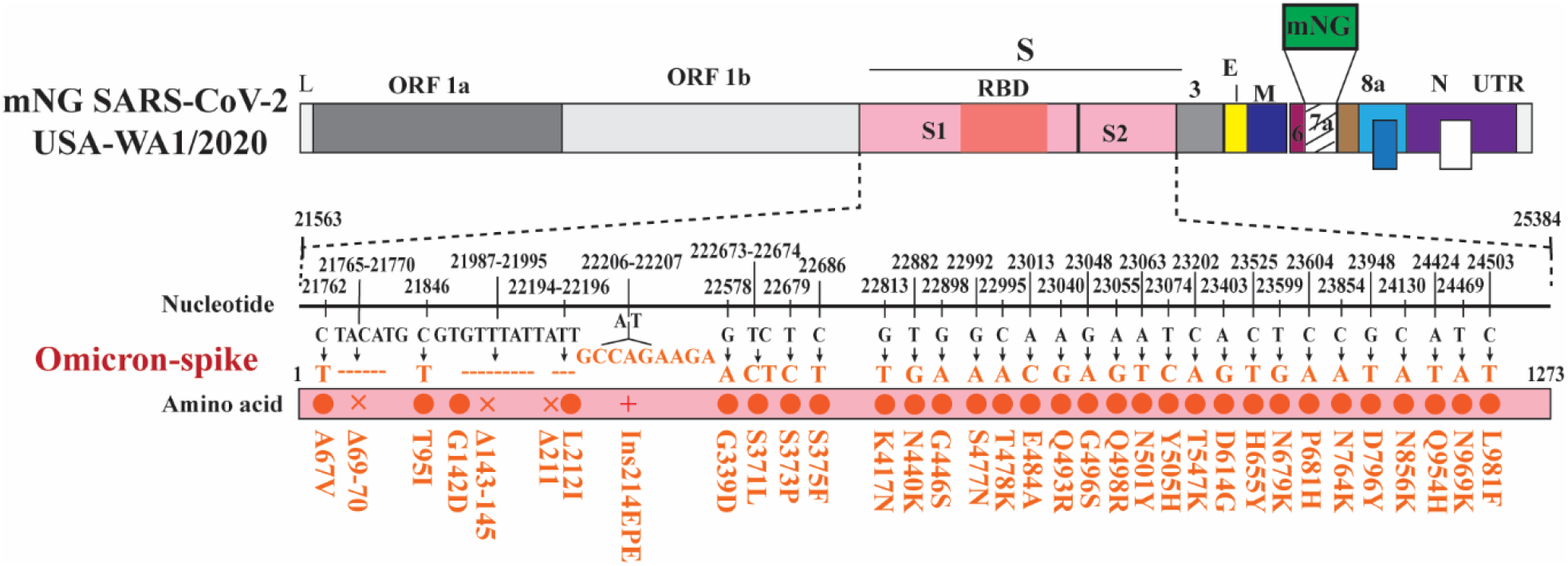
Construction of mNeonGreen (mNG) Omicron-spike SARS-CoV-2. mNG USA-WA1/2020 was used to engineer the complete *spike* gene from the Omicron variant, resulting in mNG Omicron-spike SARS-CoV-2. Mutations (red circle), deletions (x), and insertions (+) are indicated. Nucleotide and amino acid positions are depicted. L: leader sequence; ORF: open reading frame; RBD: receptor binding domain; S: spike glycoprotein; S1: N-terminal furin cleavage fragment of S; S2: C-terminal furin cleavage fragment of S; E: envelope protein; M: membrane protein; N: nucleoprotein; UTR: untranslated region.

**Extended Data Figure 2.**
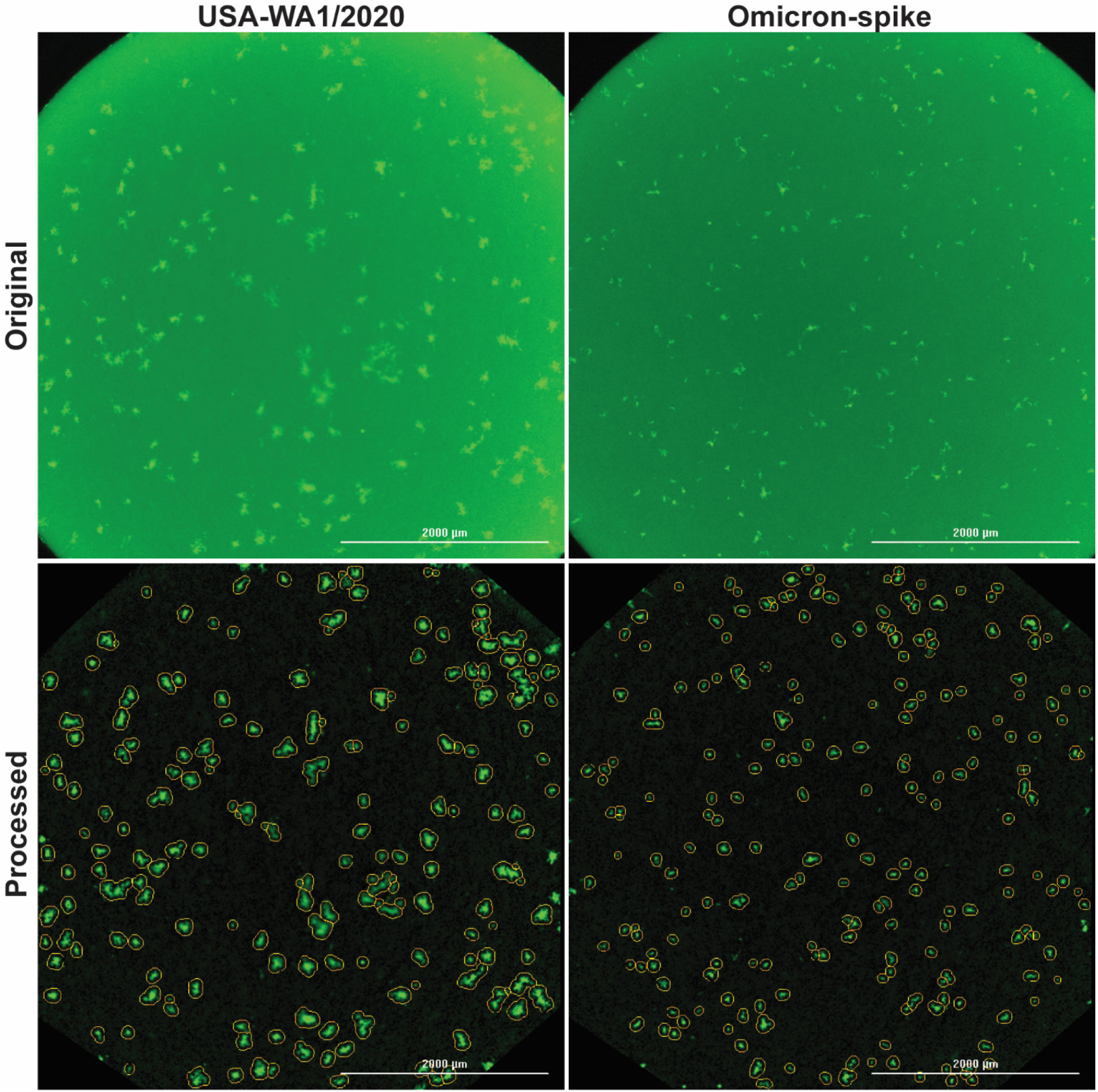
Fluorescent foci of mNG USA-WA1/2020 and mNG Omicron-spike SARS-CoV-2 on Vero E6 cells. Original and processed images were collected by high-content imaging. The protocol of the fluorescent focus reduction neutralization test (FFRNT) is described in Methods. See **Extended Data Figure 3** for the experimental scheme of FFRNT.

**Extended Data Figure 3.**
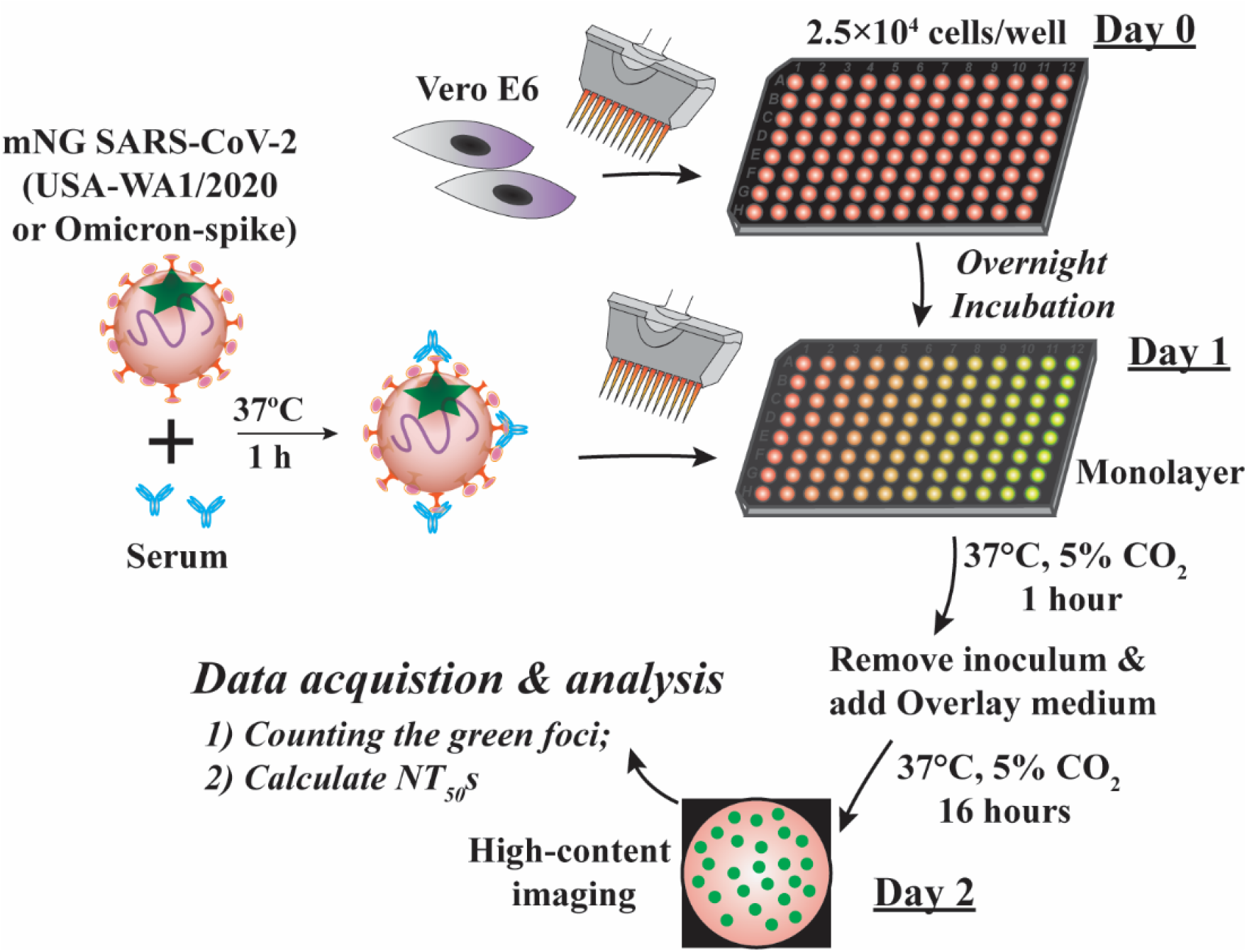
Experimental scheme of fluorescent focus reduction neutralization test (FFRNT). The FFRNT protocol is described in Methods.

**Extended Data Figure 4.**
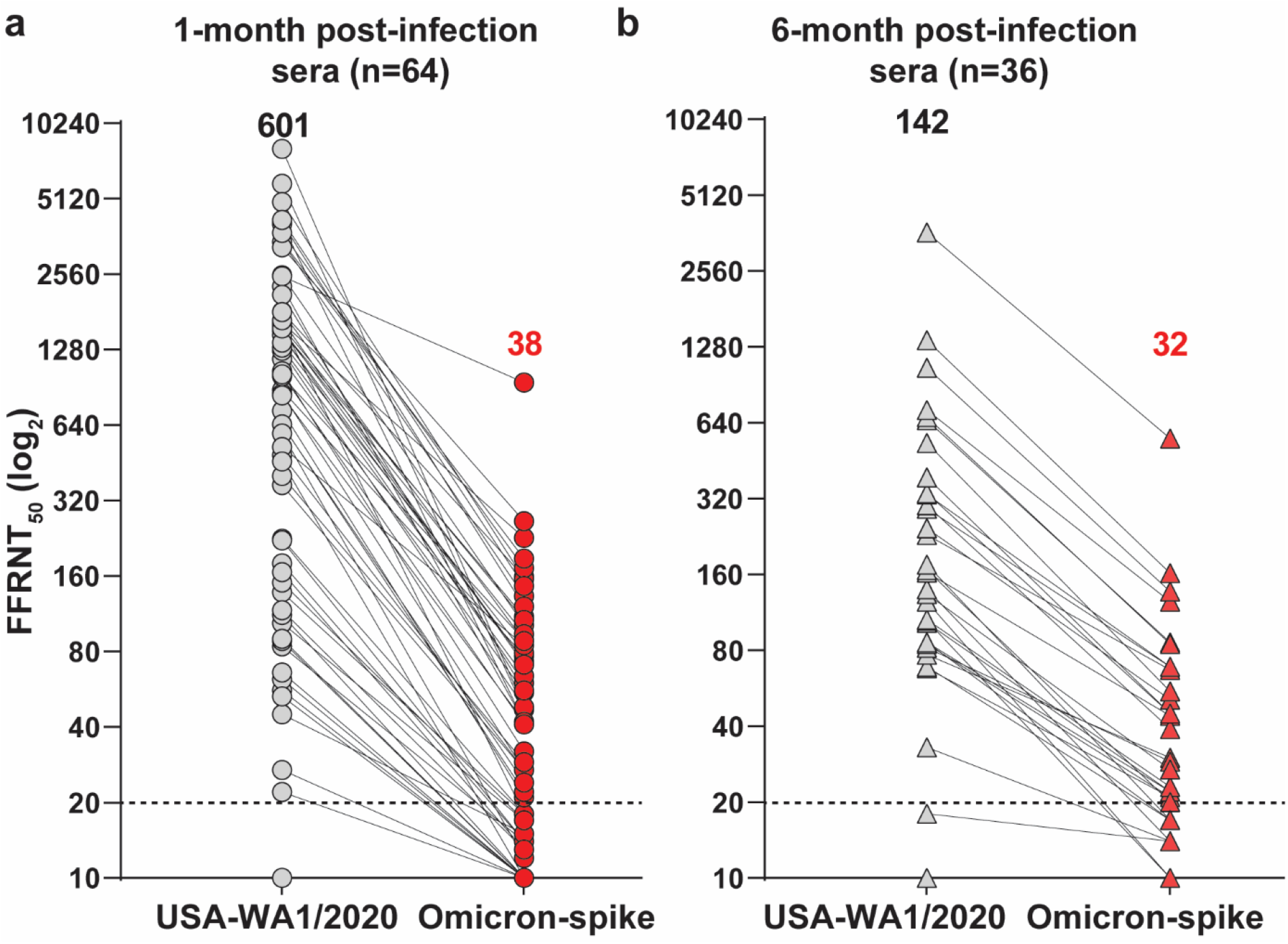
Reduced neutralization against Omicron SARS-CoV-2 by previous non-Omicron viral infection. 50% fluorescent focus reduction neutralization titers (FFRNT_50_) were measured for two serum panels from COVID-19 patients previously infected with non-Omicron SARS-CoV-2. The first serum panel was collected at 1-month post-infection (n=64) and the second panel collected at 6-month post-infection (n=36). For each serum, two FFRNT_50_ values against mNG USA-WA1/2020 and Omicron-spike SARS-CoV-2 are connected by a line. **a,** FFRNT_50_s of 1-month post-infection sera. **b,** FFRNT_50_s of 6-month post-infection sera. **Extended Data Tables 1** and **2** summarize the FFRNT_50_ values and serum information for (**a**) and (**b**), respectively. This figure is a reformat of **Figure 1**.

**Extended Data Figure 5.**
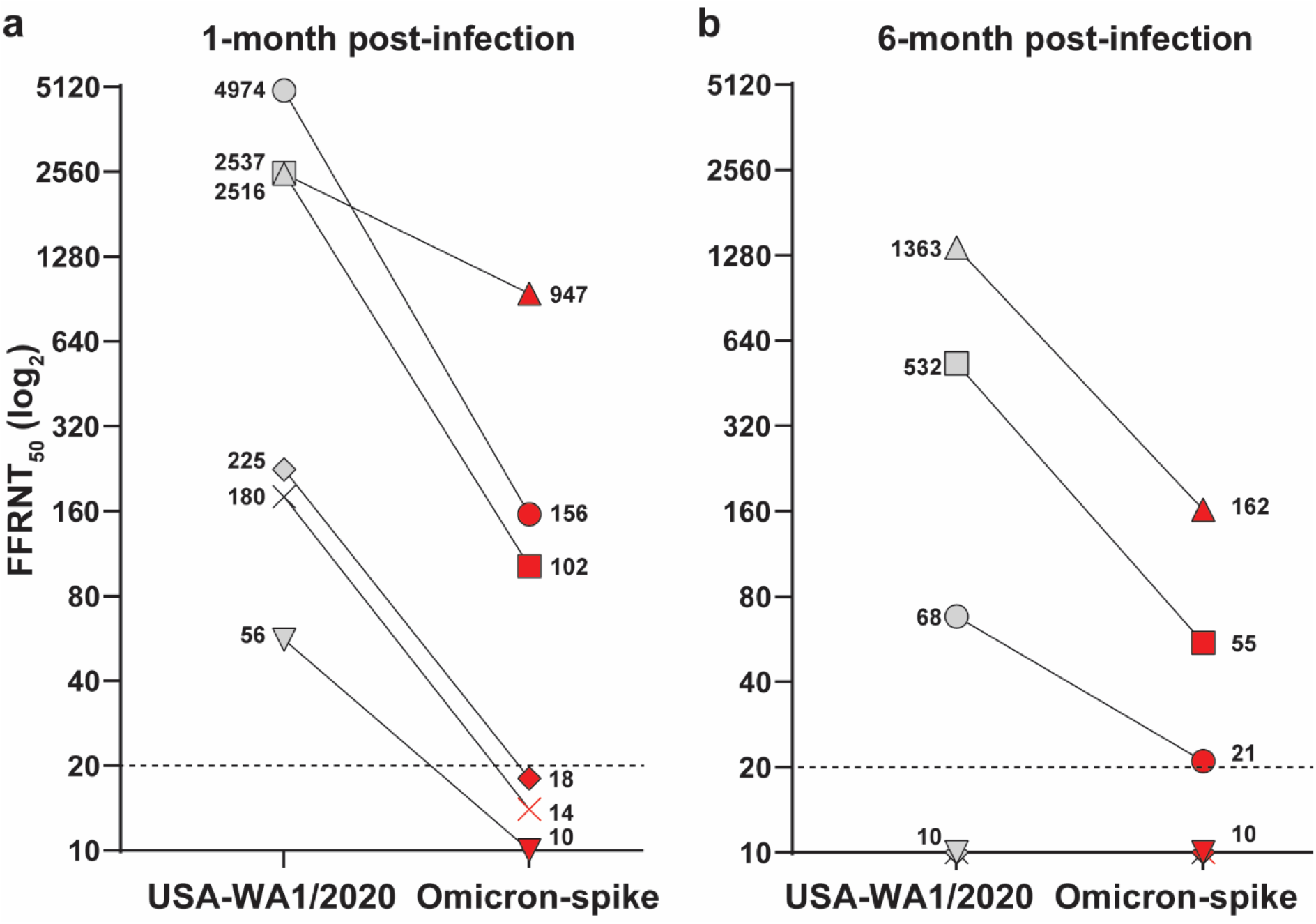
FFRNT_50_ of 6 pairs of 1- and 6-month post-infection sera from same patients. **a,** FFRNT_50_s of 1-month post-infection sera against mNG USA-WA1/2020 and Omicron-spike SARS-CoV-2. **b,** FFRNT_50_s of 6-month post-infection sera against mNG USA-WA1/2020 and Omicron-spike SARS-CoV-2.

**Extended Data Table 1.** FFRNT_50_ values of 1-month post-infection sera against mNG USA-WA1/2020 and Omicron-spike SARS-CoV-2

| Serum ID | Age | Gender | Race and Ethnicity | Sample collection date yielding positive viral test | Symptomatic | Hospitalized | FFRNT<sub>50</sub> |  |
| --- | --- | --- | --- | --- | --- | --- | --- | --- |
|  |  |  |  |  |  |  | USA-WA1/2020 | Omicron-spike |
| 1 | 21 | F | Hispanic or Latino | 7/22/2020 | No | No | 10 | 10 |
| 2 | 38 | F | White | 11/27/2020 | No | No | 22 | 10 |
| 3 | 17 | M | Hispanic or Latino | 6/1/2020 | Yes | No | 27 | 10 |
| 4 | 18 | F | Hispanic or Latino | 7/11/2020 | Yes | No | 45 | 15 |
| 5 | 26 | F | Hispanic or Latino | 11/11/2020 | Yes | No | 53 | 10 |
| 6 | 24 | F | Hispanic or Latino | 6/24/2020 | No | No | 56 | 10 |
| 7 | 24 | F | Hispanic or Latino | 7/27/2020 | Yes | No | 62 | 10 |
| 8 | 35 | F | Hispanic or Latino | 1/5/2021 | No | No | 66 | 10 |
| 9 | 23 | F | Hispanic or Latino | 6/25/2020 | No | No | 84 | 14 |
| 10 | 24 | F | Hispanic or Latino | 7/2/2020 | Yes | No | 88 | 10 |
| 11 | 33 | F | Black or African American | 11/21/2020 | Yes | No | 90 | 10 |
| 12 | 26 | F | Hispanic or Latino | 7/2/2020 | Yes | No | 104 | 17 |
| 13 | 60 | M | White | 9/26/2020 | Yes | Yes | 112 | 10 |
| 14 | 67 | M | Caucasian/White | 11/16/2020 | Yes | No | 117 | 13 |
| 15 | 22 | M | Hispanic or Latino | 9/24/2020 | No | No | 138 | 18 |
| 16 | 69 | M | White | 12/28/2020 | Yes | No | 151 | 17 |
| 17 | 73 | M | White | 5/27/2020 | Yes | No | 166 | 13 |
| 18 | 17 | F | Hispanic or Latino | 6/22/2020 | Yes | No | 180 | 14 |
| 19 | 45 | M | Hispanic or Latino | 5/2/2020 | Yes | Yes | 222 | 14 |
| 20 | 24 | F | Hispanic or Latino | 6/24/2020 | Yes | No | 225 | 18 |
| 21 | 25 | F | Black or African American | 7/16/2020 | No | No | 369 | 27 |
| 22 | 75 | F | Hispanic or Latino | 9/14/2020 | Yes | No | 402 | 29 |
| *23 | 80 | F | White | 11/9/2020 | Yes | Yes | 459 | 10 |
| 24 | 61 | F | White | 11/3/2020 | Yes | No | 486 | 27 |
| 25 | 78 | M | White | 12/15/2020 | Yes | No | 524 | 85 |
| 26 | 66 | M | White | 5/30/2020 | Yes | Yes | 592 | 12 |
| 27 | 38 | M | Hispanic or Latino | 6/27/2020 | No | No | 640 | 24 |
| 28 | 34 | M | Hispanic or Latino | 11/11/2020 | Yes | No | 645 | 22 |
| 29 | 25 | F | Hispanic or Latino | 7/17/2020 | No | No | 730 | 21 |
| 30 | 66 | M | Black or African American | 4/27/2020 | Yes | Yes | 840 | 42 |
| 31 | 39 | M | Black or African American | 4/9/2020 | Yes | Yes | 856 | 27 |
| 32 | 35 | F | Hispanic or Latino | 10/15/2020 | Yes | Yes | 882 | 77 |
| 33 | 55 | M | Hispanic or Latino | 5/6/2020 | Yes | Yes | 1003 | 89 |
| *34 | 67 | F | Hispanic or Latino | 1/4/2021 | Yes | Yes | 1020 | 41 |
| 35 | 55 | F | White | 9/28/2020 | Yes | No | 1050 | 32 |
| 36 | 40 | F | Hispanic or Latino | 12/20/2020 | Yes | No | 1174 | 55 |
| 37 | 60 | F | Hispanic or Latino | 11/27/2020 | Yes | Yes | 1268 | 48 |
| 38 | 65 | M | Hispanic or Latino | 5/7/2020 | Yes | Yes | 1306 | 64 |
| *39 | 69 | M | White | 11/22/2020 | Yes | Yes | 1365 | 79 |
| 40 | 68 | M | Caucasian/White | 5/10/2020 | Yes | Yes | 1454 | 73 |
| 41 | 50 | M | Black or African American | 4/8/2020 | Yes | Yes | 1465 | 60 |
| 42 | 63 | M | Hispanic or Latino | 1/23/2021 | Yes | Yes | 1517 | 94 |
| 43 | 39 | M | Black or African American | 3/31/2020 | Yes | Yes | 1519 | 110 |
| 44 | 72 | M | White | 12/23/2020 | Yes | Yes | 1555 | 73 |
| 45 | 55 | F | White | 10/6/2020 | Yes | Yes | 1584 | 107 |
| 46 | 57 | M | White | 7/5/2020 | Yes | No | 1618 | 18 |
| 47 | 1 | F | Hispanic or Latino | 1/18/2021 | Yes | Yes | 1638 | 227 |
| 48 | 87 | M | White | 1/5/2021 | Yes | Yes | 1679 | 71 |
| 49 | 96 | F | White | 12/30/2020 | Yes | Yes | 1807 | 88 |
| 50 | 66 | M | Hispanic or Latino | 12/19/2020 | Yes | No | 1814 | 159 |
| 51 | 75 | M | Hispanic or Latino | 10/27/2020 | Yes | No | 2119 | 56 |
| 52 | 63 | F | Hispanic or Latino | 12/12/2020 | Yes | Yes | 2289 | 42 |
| △53 | 66 | M | White | 12/27/2020 | Yes | Yes | 2516 | 947 |
| □54 | 49 | M | Black or African American | 1/3/2021 | No | Yes | 2537 | 102 |
| 55 | 56 | M | Hispanic or Latino | 7/13/2020 | Yes | Yes | 3277 | 265 |
| 56 | 44 | F | Black or African American | 8/20/20/ | Yes | Yes | 3443 | 133 |
| *57 | 83 | M | Hispanic or Latino | 11/22/2020 | Yes | Yes | 3464 | 188 |
| *58 | 75 | M | White | 12/27/2020 | Yes | Yes | 3741 | 172 |
| *59 | 74 | M | White | 12/21/2020 | Yes | Yes | 4055 | 42 |
| 60 | 48 | F | Hispanic or Latino | 6/21/2020 | Yes | Yes | 4116 | 121 |
| 61 | 78 | F | Hispanic or Latino | 12/16/2020 | Yes | Yes | 4216 | 146 |
| ○*62 | 70 | M | Hispanic or Latino | 12/12/2020 | Yes | Yes | 4974 | 156 |
| *63 | 49 | M | White | 12/29/2020 | Yes | Yes | 5876 | 47 |
| 64 | 50 | F | Hispanic or Latino | 11/9/2020 | Yes | Yes | 8088 | 48 |
| GMT | 44 | - | - | - | - | - | 601 | 38 |
| 95%CI | 37-52 | - | - | - | - | - | 405-891 | 29-50 |

\* Patients received convalescent plasma treatment.
▽×◇△□○ Patients who gave both 1- and 6-month post-infection sera.

**Extended Data Table 2.**
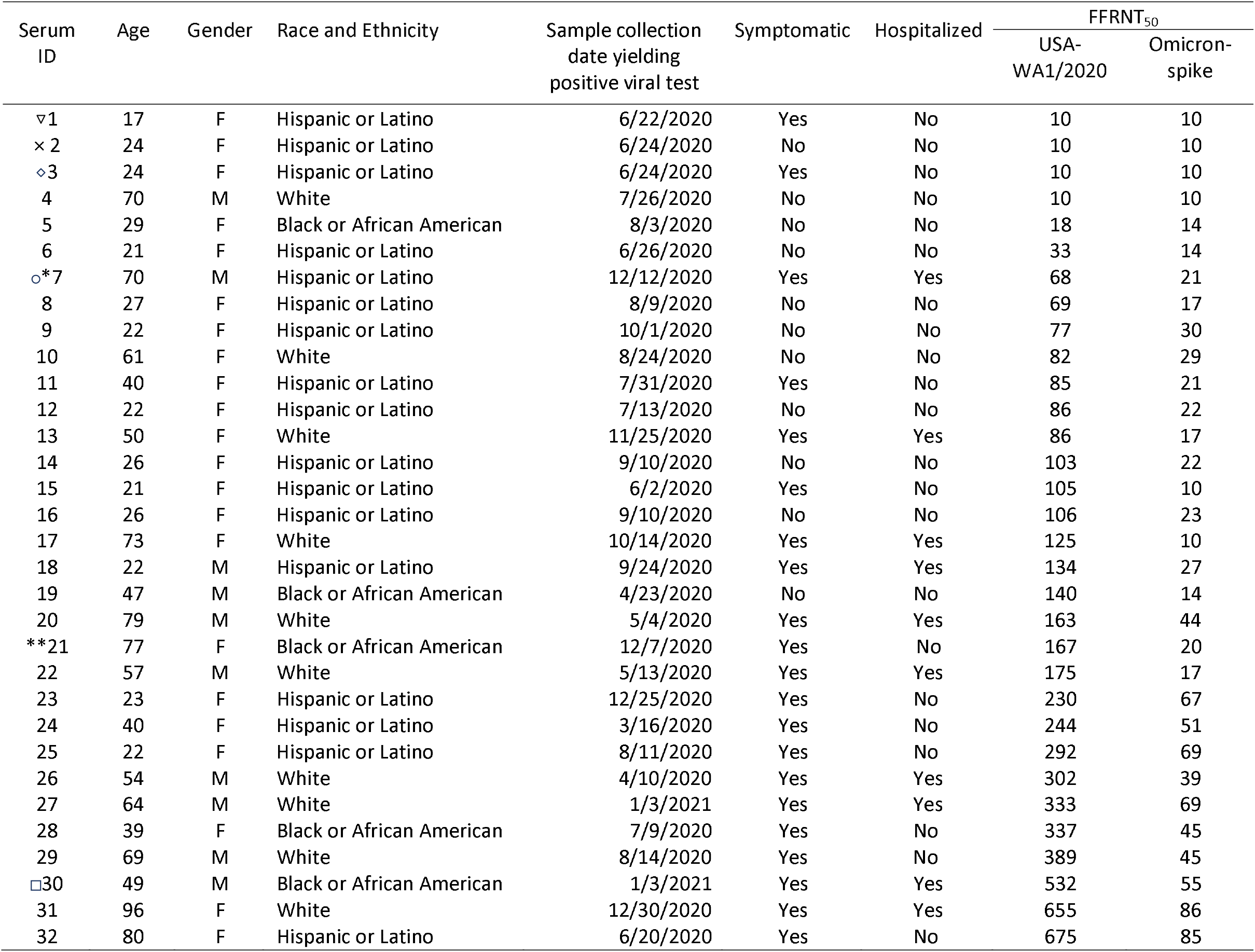
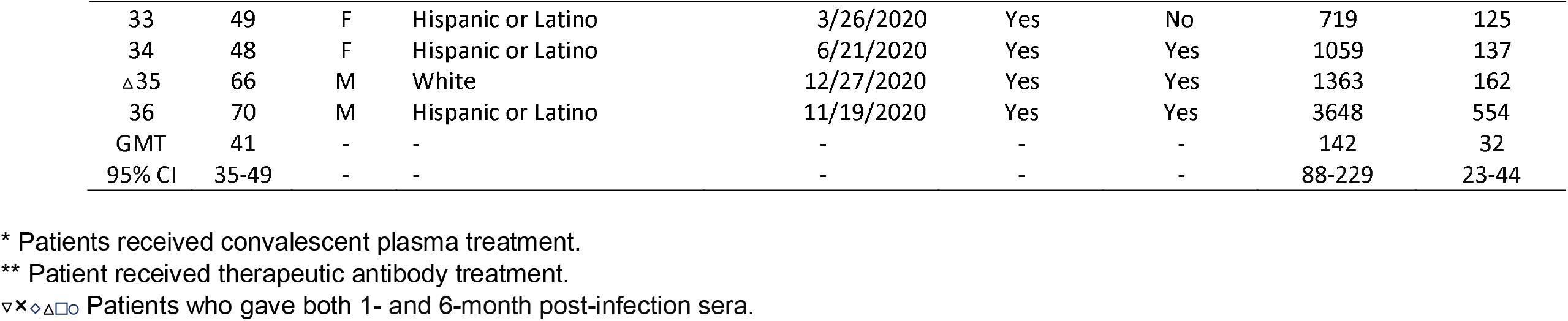
FFRNT_50_ values of 6-month post-infection sera against mNG USA-WA1/2020 and Omicron-spike SARS-CoV-2

